# The multidrug resistance efflux pump MexCD-OprJ is a switcher of the *Pseudomonas aeruginosa* quorum sensing response

**DOI:** 10.1101/322792

**Authors:** Manuel Alcalde-Rico, Jorge Olivares-Pacheco, Carolina Alvarez-Ortega, Miguel Cámara, José Luis Martínez

## Abstract

Most antibiotic resistance genes acquired by human pathogens originate from environmental microorganisms. Therefore, understanding the additional functions of these genes, other than conferring antibiotic resistance, is relevant from an ecological point of view. We examined the effect that overexpression of the MexCD-OprJ multidrug efflux pump has in the physiology of the environmental opportunistic pathogen *Pseudomonas aeruginosa*. Overexpression of this intrinsic resistance determinant shuts down the *P. aeruginosa* quorum sensing (QS) response. Impaired QS response is due to the extrusion of 4-hydroxy-2-heptylquinoline (HHQ), the precursor of the *Pseudomonas* Quinolone Signal (PQS), leading to low PQS intracellular levels and reduced production of QS signal molecules. The *P. aeruginosa* QS response induces the expression of hundreds of genes, which can be costly unless such activation becomes beneficial for the bacterial population. While it is known that the QS response is modulated by population density, information on additional signals/cues that may alert the cells about the benefits of mounting the response is still scarce. It is possible that MexCD-OprJ plays a role in this particular aspect; our results indicate that, upon overexpression, MexCD-OprJ can act as a switcher in the QS population response. If MexCD-OprJ alleviate the cost associated to trigger the QS response when un-needed, it could be possible that MexCD-OprJ overproducer strains might be eventually selected even in the absence of antibiotic selective pressure, acting as antibiotic resistant cheaters in heterogeneous *P. aeruginosa* populations. This possibility may have potential implications for the treatment of *P. aeruginosa* chronic infections.

## Importance

It has been proposed that antibiotic resistance genes might have ecological functions going beyond antibiotic resistance. The role that the *Pseudomonas aeruginosa* multidrug efflux pump MexCD-OprJ may have in intercellular signaling was explored. Overexpression of this resistance determinant shuts down the *P. aeruginosa* quorum sensing (QS) response via extrusion of QS signals/precursors. A function of this efflux pump, and of others that reduce the QS response when overexpressed, could be acting as a QS switch of *P. aeruginosa* in response to environmental cues, allowing to switch-off the system in those conditions in which the activation of the energy-expensive QS response is not advantageous, despite the population density being high enough to trigger such response. MexCD-OprJ overproducers might be eventually selected even in the absence of antibiotic selective pressure, acting as antibiotic resistant cheaters in heterogeneous *P. aeruginosa* populations, which has potential implications for the treatment of *P. aeruginosa* infections.

## Introduction

*Pseudomonas aeruginosa* is a free-living microorganism able to survive in different environments that not only plays an ecological role in natural ecosystems (1–4), but it is also an important causative agent of infections in patients with underlying diseases (5–8). The characteristic low susceptibility to antibiotics of this organism relays on several factors (9). Particularly relevant is the activity of chromosomally-encoded multidrug-resistance (MDR) efflux pumps (10, 11). Further, the acquisition of mutation-driven resistance is common in this opportunistic pathogen, particularly along chronic infections (12–14), where the constitutive overexpression of MDR efflux pumps is one of the biggest problems to eradicate these infections (10, 15, 16). Efflux pumps exhibit different functions, with physiological and ecological significances that go beyond their activity as antibiotic resistance elements (10, 17–19). In the case of *P. aeruginosa*, an opportunistic pathogen not fully adapted to human hosts (1, 20), these functions should be of relevance for the success of *P. aeruginosa* to colonize natural ecosystems.

*P. aeruginosa* harbours several efflux systems that belong to different families (21). The most studied because of their clinical relevance are MexAB-OprM (22), MexCD-OprJ (23, 24), MexEF-OprN (24), and MexXY (25, 26). They all belong to the *Resistance-Nodulation-Division* (RND) family of MDR systems (10). Mutants that exhibit constitutive overexpression of each of these efflux pumps are selected upon treatment with antibiotics; the mutations are frequently located in the regulatory elements adjacent to the respective operon encoding for these MDR systems (27–30).

The unregulated overexpression of an efflux system not only contributes to antibiotic resistance but may also have pleiotropic effects in the bacterial physiology. We have recently reported that overexpression of RND systems in *P. aeruginosa* leads to an excessive internalization of protons that acidify the cytoplasm, which causes a biological cost in absence of oxygen or nitrate, since both are necessary to compensate for the intracellular H^+^ accumulation (31, 32). In addition to these non-specific effects, other effects might be due to the unregulated extrusion of intracellular compounds, some of which may be relevant for the ecological behaviour of *P. aeruginosa* (33). Indeed, different studies have shown that overexpression of MDR efflux pumps may challenge the *P. aeruginosa* quorum sensing (QS) response (34–37), which is in turn determinant for modulating several physiological processes in response to population density (38).

In *P. aeruginosa*, the QS-signalling network consists of three main interconnected regulatory systems: Las, Rhl, and Pqs, which synthetize and respond to the autoinducers *N*-(3-oxododecanoyl)-L-homoserine lactone (3-oxo-C12-HSL), *N*-butanoyl-L-homoserine lactone (C4-HSL), and the 2-alkyl-4(1*H*)-quinolones (AQs) *Pseudomonas* Quinolone Signal (PQS, or its immediate precursor 2-heptyl-4-hydroxyquinoline, HHQ), respectively (39). These autoinducers are able to bind to their respective transcriptional regulators, namely LasR, RhlR and PqsR, thus controlling the expression of a large number of genes including those responsible for their own synthesis: *lasI*, *rhlI* and *pqsABCDE* respectively.

Some *P. aeruginosa* RND systems have been associated with QS. MexAB-OprM is induced by C4-HSL (40) and has been proposed to extrude 3-oxo-C12-HSL and other 3-oxo-HSL related compounds (36, 41, 42). MexEF-OprN is able to efflux HHQ (43) and kynurenine (34), both precursors of the PQS autoinducer signal (44, 45). In agreement with these findings, the antibiotic resistant mutants that overproduce MexAB-OprM or MexEF-OprN have been associated with a low production of QS-controlled virulence factors (34, 36, 42, 46). Some studies have demonstrated that acquisition of antibiotic resistance due to constitutive overexpression of *mexCD-oprJ* correlates with a decrease in the production of several virulence factors, some of them controlled by QS (28, 35, 46, 47). However, the underlying reasons for this correlation remain to be elucidated. In this work, we analysed in depth the production of each QS signal molecule (QSSM) and the expression levels of the genes controlled by these regulation systems in order to understand how overexpression of MexCD-OprJ could be affecting the *P. aeruginosa* QS response.

## Results

Increased expression of efflux pumps due to mutations in their regulators can produce different changes in bacterial physiology. In most cases, the phenotypes observed in this kind of mutants are attributed to the activity of the overexpressed efflux pump. However, in other instances, the mutations in the local regulator itself might have effects on the bacterial physiology independently of the activity of the efflux pump (48, 49). To address this possibility, we used a previously described mutant that overexpresses MexCD-OprJ (35). To discard the possibility that other mutations besides those in the *mexCD-oprJ* repressors might have been selected in this strain during its stay in the laboratory, the genome of the mutant was fully sequenced. Only the already described *nfxB* mutation (35) was found. From this mutant, an *nfXB*\**ΔmexD* strain, which keeps the *nfxB* mutation in addition to a partial deletion of the *mexD* gene, was generated. By comparing *nfxB*\* and *nfXB*\**ΔmexD* strains, we were able to define more precisely which phenotypes depend on the activity of the efflux pump and which are solely due to the inactivation of the NfxB repressor, independently of the activity of the efflux pump.

### Overexpression of MexCD-OprJ results in a decrease in the production of QS-controlled virulence factors in *P. aeruginosa*

Swarming motility and the production of elastase, proteinase IV, pyocyanin, and, rhamnolipids were analysed to establish whether or not MexCD-oprJ affects the production of *P. aeruginosa* QS-regulated virulence elements. As Figure 1 shows and in agreement with previous studies (46, 47), the *nfxB*\* strain exhibits a decrease in swarming motility and in the production of all analysed virulence factors in comparison with the wild-type PAO1 strain. The fact that the deletion of *mexD* fully restores the production of QS-regulated virulence factors in an *nfxB*\* background, indicates that the observed impairment is solely due to the activity of MexCD-OprJ, independently of the potential activity of the NfxB regulator protein.

**Figure 1.**
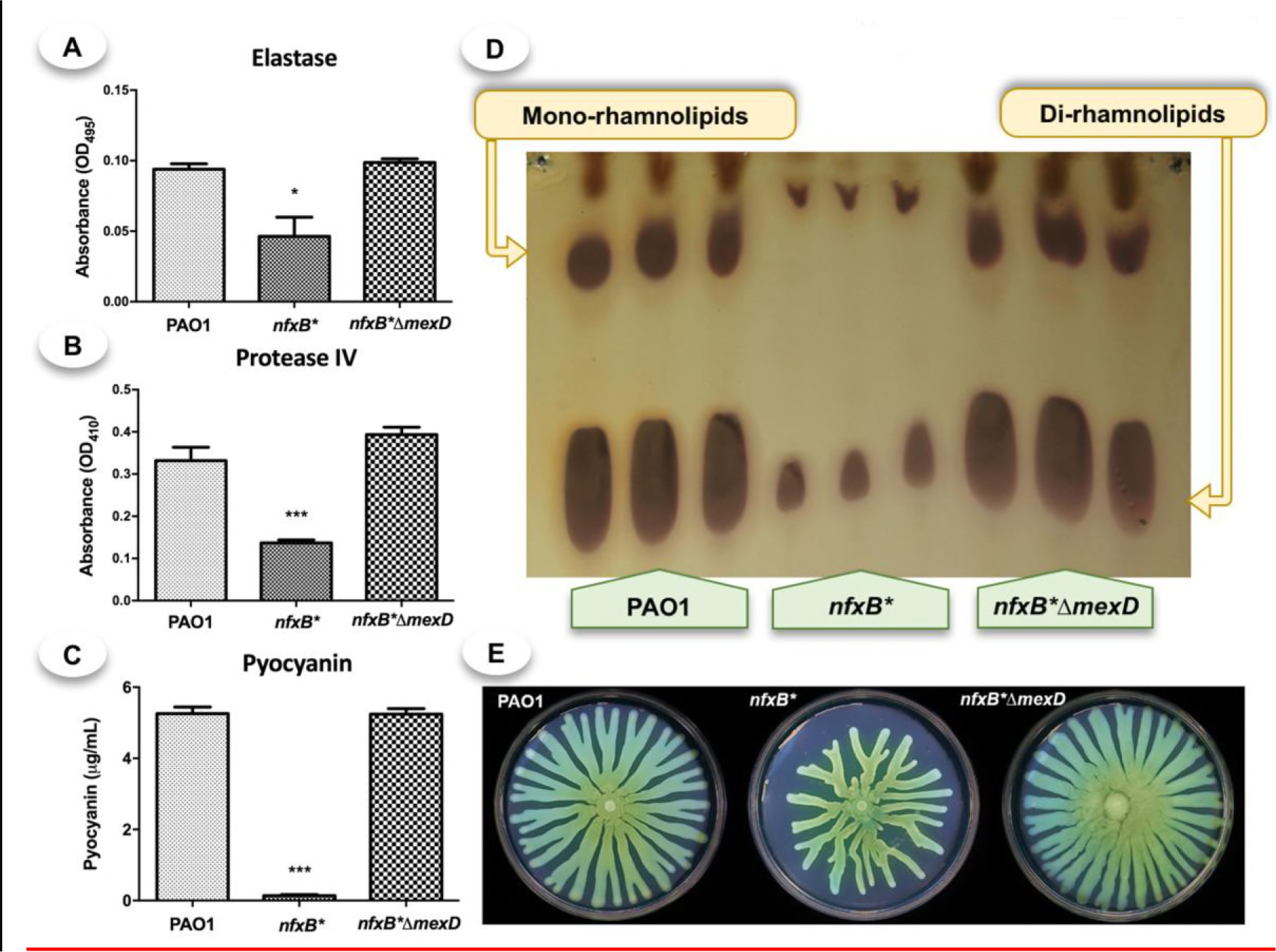
Overexpression of the MexCD-OprJ efflux pump results in a decrease in the production of different virulence factors regulated by the QS system. The elastase (A), protease IV (B), pyocyanin (C) and rhamnolipids (D) assays were conducted with supernatants of LB liquid cultures of the PAO1, *nfxB*\* and *nfXB*\**ΔmexD* strains after 20 hours of incubation at 37° C. *nfxB*\* presented a lower production of all tested virulence factors that the parental wild-type PAO1. The deletion of *mexD* in strain *nfXB*\**ΔmexD* restores the phenotypes to the levels of the wild-type strain, indicating that the defects in the expression of virulence factors were solely due to the activity of the *mexCD-oprJ*_efflux pump.

### Overproduction of the MexCD-OprJ efflux system results in a lower expression of QS- regulated genes

Expression of a set of QS-regulated genes (50–54) was analysed to determine if a low production of virulence factors in the *nfxB*\* mutant correlates with a deregulated expression of QS-regulated genes. LasB controls elastase production (52, 55). RhlA and RhlB are implicated in rhamnolipids biosynthesis (52, 53), which in turn is important for swarming motility (53). PhzB1, PhzB2, and PhzS are implicated in pyocyanin biosynthesis and the MexGHI-OpmD efflux pump has been described to be regulated by this phenazine (54). As shown in Figure 2A, the expression levels of the tested genes are lower in the *nfxB*\* strain than in PAO1. In addition, the expression of these genes is restored to PAO1 levels upon *mexD* deletion in the *nfxB*\* strain, further confirming that *mexCD-oprJ* overexpression of is what causes an impaired QS response in the *nfxB*\* mutant. These results are in agreement with the lower production of virulence factors observed in *nfxB*\* (Figure 1).

**Figure 2.**
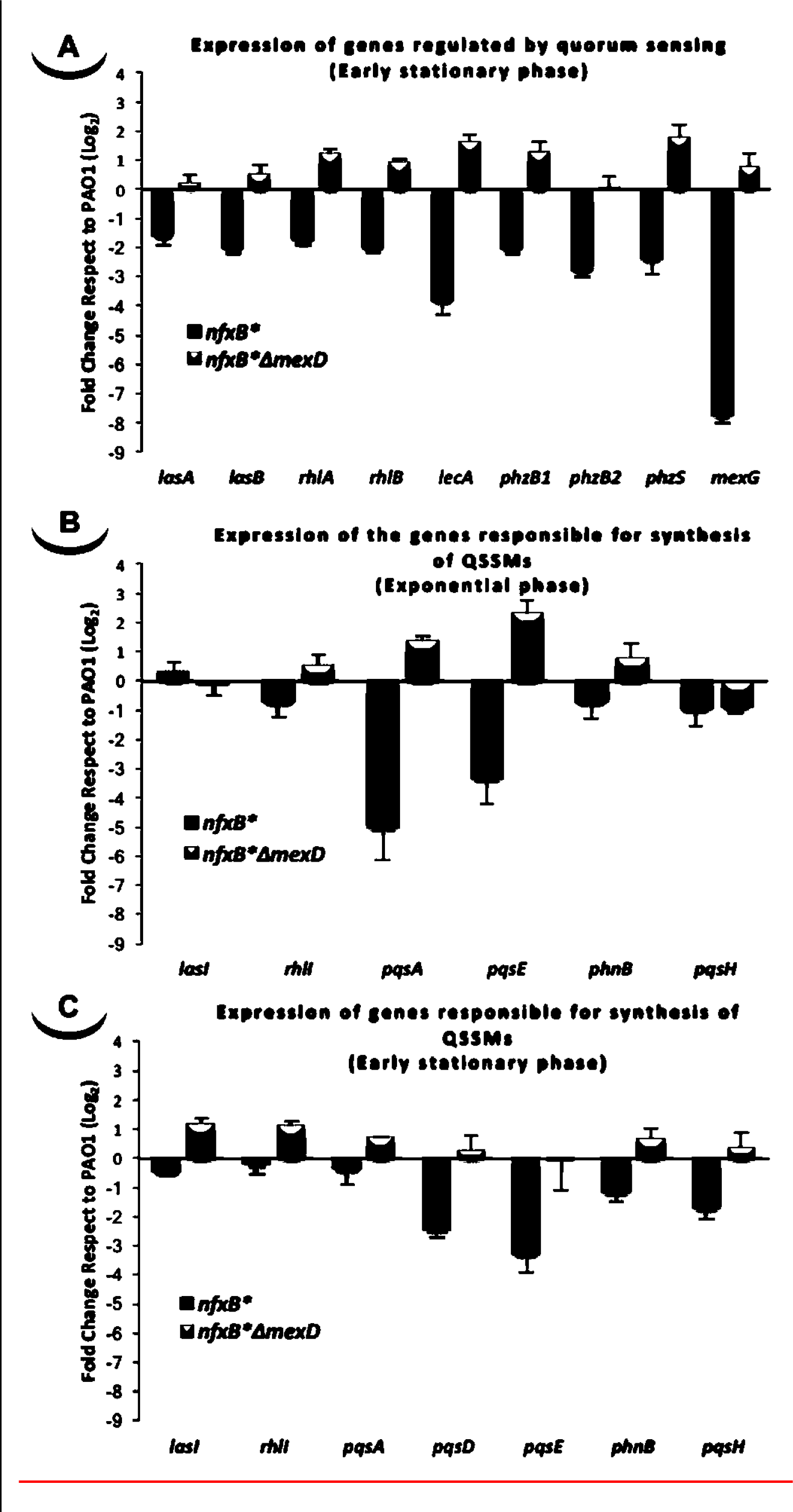
Overexpression of the MexCD-OprJ efflux system affects the expression levels of QS-regulated genes. Transcriptional analysis by real-time RT-PCR of (A) genes regulated by quorum sensing response (*lasA*, *lasB*, *rhlA*, *rhlB*, *lecA*, *phzB1*, *phzB2*, *phzS* and *mexG*) and (B and C) genes responsible for QSSMs production (*lasI*, *rhlI*, *pqsA*, *pqsD*, *pqsE*, *phnB* and *pqsH*) from samples obtained in (B) exponential (OD_600_ = 0.6) and (A and C) early stationary phase of growth (OD_600_ = 2.5) in PAO1, *nfxB*\* and *njXB*ΔmexD* strains grown in LB medium. The results showed that the gene responsible for C4-HSL production (*rhlI*) and some genes implicated in the synthesis of PQS (*pqsA, pqsE*, and *phnB*) were expressed at lower level in the *nfxB*\* strain than in the wild-type PAO1 strain at exponential phase of growth (B). In early stationary growth phase (A and C), the expression levels of the PQS-biosynthesis genes (*pqsD*, *pqsE, phnB* and *pqsH*), as well as all of the analysed QS-regulated genes, were much lower in the *nfxB*\* strain than in the PAO1 wild-type. The deletion of *mexD* in strain *nfXB*\**ΔmexD* restores the levels of expression to those of the wild-type strain, indicating that these defects were solely due to the activity of the MexCD-OprJ_efflux pump.

To gain more insights on the reasons for this impaired QS-response, we analysed the expression of genes responsible for the production of both families of autoinducers AHLs (*lasI* and *rhlI*) (56) and AQs (*pqsABCDE-phnAB* and *pqsH*) (57). This was performed along the exponential growth phase when expression of these QS biosynthesis genes starts, and in early stationary phase, when the Pqs-system is fully active (58). As shown in Figures 2B and 2C, expression of the genes responsible for the synthesis of PQS and HHQ exhibit a marked decrease in the *nfxB*\* strain at both time points. These changes were restored to wild-type levels upon MexCD-OprJ inactivation in an *nfxB*\* background. *pqsA*, from the *pqsABCDE* operon responsible for the biosynthesis of AQs (57), exhibits the sharpest decrease in expression during exponential growth phase (Figure 2B). Expression of *phnB*, implicated in the synthesis of anthranilate through the chorismic acid pathway (44, 45), as well as *pqsH*, which codify the enzyme responsible for the conversion of HHQ into PQS (57), decreases more in early stationary phase (Figures 2B and 2C).

In contrast to the strong variations in expression of PQS-related genes, the activity of MexCD-OprJ had a minor impact on the expression of AHLs-related genes in both exponential and stationary growth phases. The *nfxB*\* strain did not present alterations in *lasI* expression, the gene responsible for the synthesis of 3-oxo-C12-HSL, neither in exponential (Figure 2B) nor in stationary phase of growth (Figure 2C). A similar behaviour was observed for *rhlI*, detecting just a slight decreased expression in the *nfxB*\* strain during exponential growth phase (Figures 2B and 2C).

### MexCD-OprJ overexpression entails a decrease in the production and accumulation of AQs due to their extrusion through this efflux pump

The production and accumulation of PQS and HHQ in both supernatant and cellular extracts decreased in the *nfxB*\* mutant (Figure 3A). This effect is directly dependent on MexCD-OprJ activity, since PQS/HHQ accumulation was restored to nearly wild-type levels in the *njXB*ΔmexD* strain (Figure 3A). Interestingly, the proportion of HHQ present in the supernatants with respect to cell-extracts is different among the three strains. As Figure 3B shows, the *nfxB*\* mutant has a higher supernatant/cell extract HHQ ratio than PAO1. Further, the deletion of *mexD* in the *nfxB*\* strain restored the HHQ ratio to similar values than those of the wild-type strain, suggesting that MexCD-OprJ may be extruding HHQ, affecting the progressive intracellular accumulation of this signal. Since the expression of the *pqsABCDE-phnAB* operon, responsible of AQs biosynthesis (57), is activated in presence of PQS/HHQ (50, 59, 60), we postulate that HHQ extrusion by MexCD-OprJ could be the main cause for the lower production of AQs observed in the *nfxB*\* strain, ultimately resulting in a defective QS-system.

**Figure 3.**
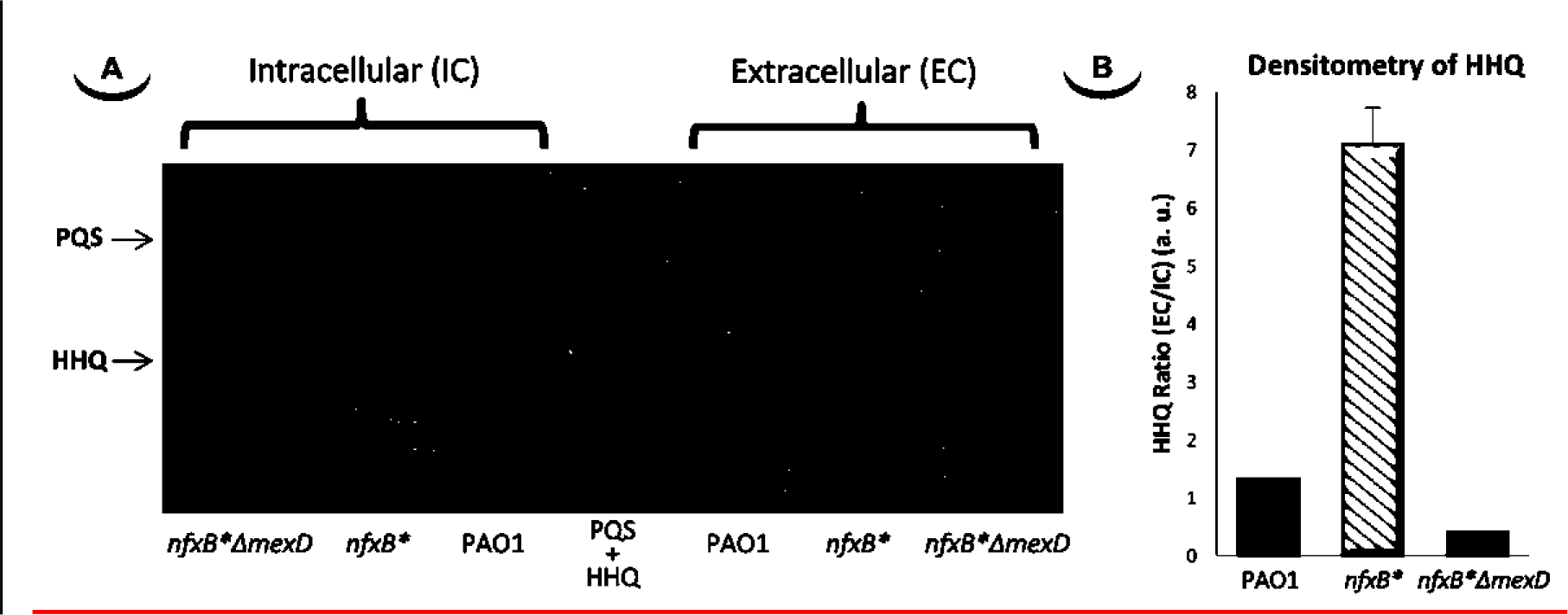
PQS and HHQ production is impaired in the strain that overproduces the MexCD-OprJ efflux pump. (A) To determine the accumulation levels of the autoinducers synthesized by *P. aeruginosa*, a technique based on TLC coupled with a PqsR-based biosensor was used. The samples were extracted from cultures in early stationary phase (OD_600_ = 2.5). (B) The TLC-spots corresponding to HHQ were quantified by densitometry and the ratio between the HHQ present in the supernatant respect to cell extract was calculated and represented. As shown, overexpression of the MexCD-OprJ efflux system in *nfxB*\* strongly reduces the production of PQS and HHQ as compared with PAO1 and *nfXB*\**ΔmexD* strains. Furthermore, the analysis by densitometry of the HHQ ratio shows that this defect in AQs production is likely caused by an excessive extrusion of HHQ through MexCD-OprJ.

### Overexpression of MexCD-OprJ produces minor effects in the synthesis of 3-oxo-C12-HSL and C4-HSL autoinducers

Since the Las, Rhl and Pqs regulation systems are highly interconnected (61–63), we wanted to know whether or not the excessive HHQ extrusion through MexCD-OprJ in the *nfxB*\* mutant could be also affecting the production of the QS signals, 3-oxo-C12-HSL (autoinducer signal for Las system) and C4-HSL (autoinducer signal for Rhl system). As shown in Figure 4, both intracellular and extracellular amounts of 3-oxo-C12-HSL are slightly higher in *nfxB*\* cultures than in either the wild-type PAO1 strain or the *njxB*ΔmexD* mutant. The opposite effect was observed for C4-HSL; the *nfxB*\* mutant accumulates slightly lower extracellular levels of this QS signal during exponential phase, reaching the levels of extracellular accumulation observed in both PAO1 and *njxB*ΔmexD* in early stationary phase (Figures 4D and 4E). This variation may also exist inside the cell due to the ability of C4-HSL to freely diffuse through cytoplasmic membrane (42). Altogether, these results indicate that overexpression of MexCD-OprJ leads to minor alterations of AHLs production. These changes might be due to the strongly impaired production of PQS and HHQ.

**Figure 4.**
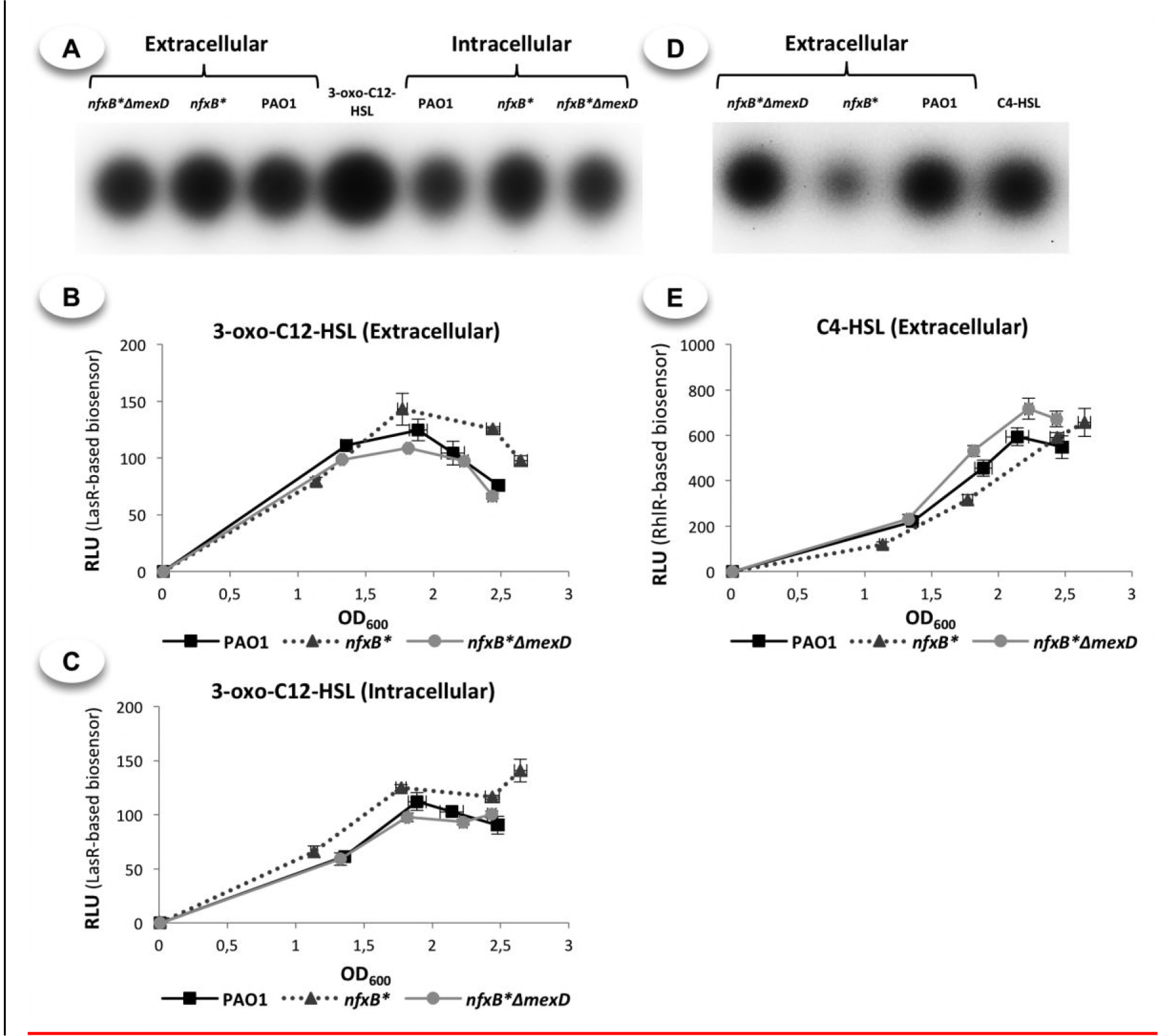
The *nfxB*\* mutant displays minor alterations in the kinetic of accumulation of both C4-HSL and the 3-oxo-C12-HSL autoinducers. TLCs (A and D) and time course accumulation assays (B, C and E) were used to determine the accumulation of these two autoinducer compounds. The samples for the TLC assays were extracted from cultures in late exponential phase (OD_600_ = 1.7) and the samples for the time course assay were taken at different time along the cell cycle (4, 5, 6 and 7 hours post-inoculation). As shown, the overexpression of MexCD-OprJ has a slightly but significant effect on AHLs accumulation. The *nfxB*\* strain presented higher levels of 3-oxo-C12-HSL than PAO1 and *njxB*ΔmexD* both outside (A and B) and inside the cells (A and C). In contrast, the C4-HSL accumulation in the supernatant was lower in the MexCD-OprJ overexpressing mutant as compared with PAO1 and *njxB*ΔmexD* strains, although similar levels were detected once the three strains reached stationary phase (6 and 7 hours after inoculation).

### MexCD-OprJ is able to extrude kynurenine but not anthranilate, both precursors of AQs signals

Our results indicate that the impaired QS response associated to the overexpression of the MexCD-OprJ efflux pump is mainly caused by a decreased production of PQS and HHQ, likely due to an excessive HHQ extrusion through this efflux system. The MexEF-OprN efflux pump is able to extrude both HHQ and its precursor kynurenine (34, 43); extrusion of the latter is the main cause for the impaired QS response observed in MexEF-OprN overproducer strains (34). A similar situation might also apply to MexCD-OprJ.

One of the immediate precursors of AQs in *P. aeruginosa* is anthranilate, which is mainly synthetized from tryptophan when this amino acid is present in the medium through the kynurenine pathway, while it is synthetized from chorismate when tryptophan is absent (Figure 5) (44, 64). Since the kynurenine pathway is the main source of anthranilate for AQs production when bacteria grow in rich LB medium (44), it could be possible that extrusion of some of the biosynthetic intermediates through MexCD-OprJ might affect the AQs production in *nfxB\**. To test this hypothesis, we first analysed the growth kinetic of PAO1 and *nfxB*\* in minimal medium containing tryptophan, kynurenine or succinate as the sole carbon source. As shown in Figure 6A, the *nfxB*\* mutant presents a growth defect in both tryptophan or kynurenine as the sole carbon source when compared to PAO1, which strongly suggests extrusion of one or more intermediates of the kynurenine pathway.

**Figure 5.**
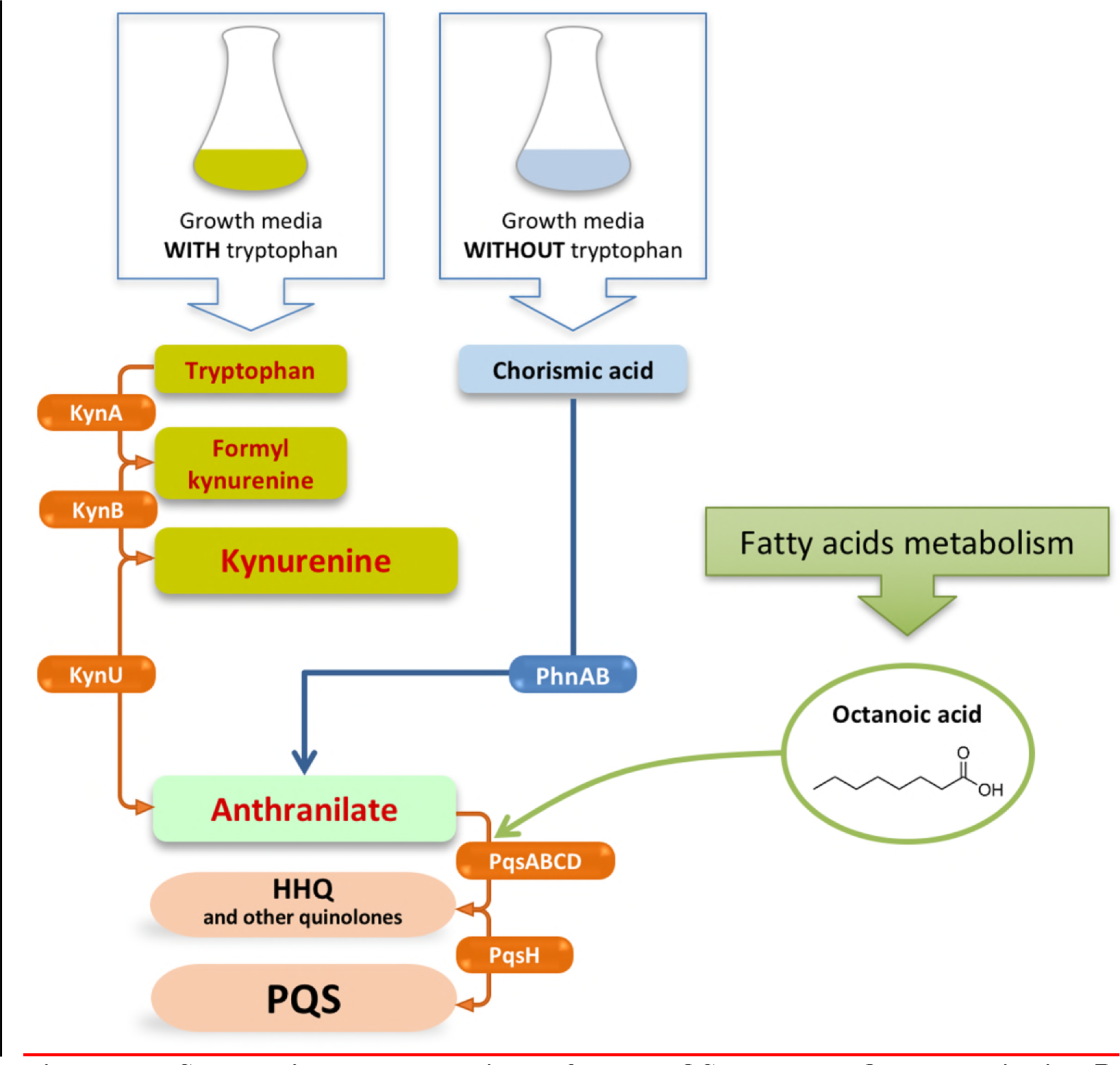
Schematic representation of the PQS and HHQ synthesis in *P. aeruginosa*. The enzymes responsible for the synthesis of HHQ and other quinolones are codified in the operon *pqsABCDE*. These enzymes use anthranilate and octanoate as precursor of HHQ and the enzyme PqsH is responsible to transform it into PQS The main source of octanoate is provided from fatty acids metabolism. In the case of anthranilate, it may be synthetized by two main pathways, for subsequent PQS production, depending of the presence/absence of tryptophan: In presence of tryptophan, the main source of anthranilate is provided from kynurenine pathway; while, in absence of this amino acid, the anthranilate is mainly synthetized from chorismic acid, being implicated the enzymes PhnA and PhnB codified in the operon *pqsABCDE-phnAB*.

To verify this possibility, we looked for the presence of kynurenine and anthranilate in the supernatants of PAO1 and *nfxB*\* cultures. We observed a lower amount of anthranilate and a higher accumulation of kynurenine in the supernatants of *nfxB*\* cultures (Figure 6B). Altogether, these results show that MexCD-OprJ is able to extrude kynurenine, but not anthranilate.

**Figure 6.**
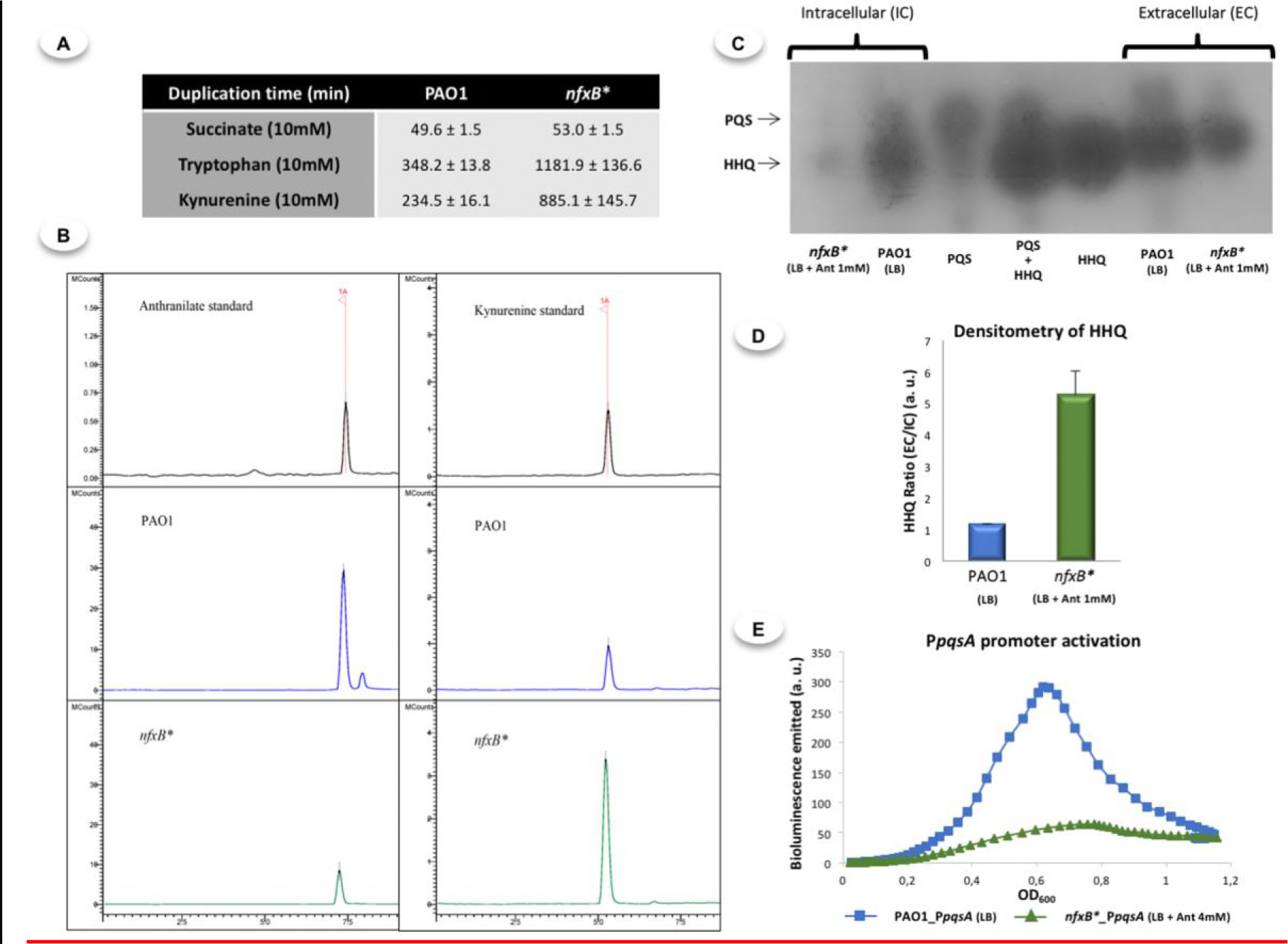
Impaired intracellular accumulation of anthranilate produced by an excessive kynurenine extrusion through MexCD-OprJ is not the cause for lower AQs production of the *nfxB*\* mutant. (A) The duplication time of PAO1 and *nfxB*\* in minimal medium with succinate (control), tryptophan or kynurenine (both anthranilate precursors) as sole carbon sources was determined. As shown, the *nfxB*\* mutant presents an impaired growth in both tryptophan and kynurenine, suggesting these compounds might be substrates of MexCD-OprJ. (B) Anthranilate and kynurenine accumulation in cell-free supernatants was quantified by HPLC-MS from PAO1 and *nfxB*\* cultures grown along 24 hours in M63 minimal medium with succinate (10 mM) and tryptophan (10 mM) as sole carbon sources. Left panels anthranilate, right panels kynurenine. As shown, the supernatants from *nfxB*\* cultures contained more kynurenine and less anthranilate than those from the wild-type PAO1 strain, indicating that kynurenine is a substrate of MexCD-OprJ and anthranilate is not extruded by this efflux pump. (C) The production of AQs in PAO1 and *nfxB*\* strains growing in LB medium supplemented with anthranilate 1 mM was analysed in early stationary phase (OD_600_ = 2.5) by TLC. (D) The extracellular vs intracellular HHQ ratios were calculated measuring each one of the HHQ spots obtained in the TLC-assays by densitometry. (E) Real-time *pqsABCDE* expression was analysed in both PAO1 and *nfxB*\* strains growing in LB medium supplemented with anthranilate 4 mM using a chromosomal insertion of the reporter construction *PpqsA::luxCDABE*. The results show that anthranilate supplementation of LB medium does not restore the AQs production in the *nfxB*\* strain (C and E), reinforcing our hypothesis that HHQ extrusion (D) rather than kynurenine extrusion through MexCD-OprJ is the main cause for the QS-defective response of the *nfxB*\* strain.

### The low levels of PQS and HHQ observed in the *nfxB*\* strain is not just due to kynurenine extrusion

Having established that the constitutive overexpression of MexCD-OprJ efflux pump leads to a decrease in the extracellular accumulation of anthranilate, we wondered whether a low intracellular availability of anthranilate could be the cause of the impaired PQS and HHQ production observed in the *nfxB*\* strain. To address this possibility, we grew PAO1 and *nfxB*\* in LB medium supplemented with 1 mM anthranilate and analysed the production and accumulation of these two signals. As shown in Figure 6C, anthranilate supplementation does not restore PQS/HHQ production to wild-type levels in the *nfxB*\* strain. In addition, our results indicate that the *nfxB*\* strain continues to extrude HHQ at higher levels than those observed in the wild-type strain under these conditions (Figure 6D).

We entertained the possibility that a higher anthranilate concentration was needed to restore AQs production to wild-type levels in *nfxB\**. To this end, we supplemented LB medium with up to 4 mM anthranilate and analysed the activation of the *pqsABCDE* promoter in real-time in both PAO1 and *nfxB\**. As shown in Figure 6E, a higher concentration of anthranilate did not restore the activation of the *pqsABCDE* promoter in the *nfxB*\* strain. These results indicate that a low anthranilate concentration caused by kynurenine extrusion is not the main underlying cause for the impaired PQS and HHQ production observed in this strain. These results further support the notion that an excessive, non-physiological, extrusion of HHQ caused by the overexpression of MexCD-OprJ is likely the main cause for the lower accumulation and production of HHQ and PQS in the multidrug resistant *nfxB*\* mutants.

### The low production of AQs associated to MexCD-OprJ overexpression is not caused by an impaired intracellular accumulation of octanoate

Octanoate is the other direct precursor of PQS and HHQ (65). Once we established that anthranilate synthesis is not the limiting step in the production of AQs by the *nfxB*\* strain, we wondered whether a hypothetical low production or intracellular accumulation of octanoate might be affecting the AQs production in this strain. For that purpose, we measured the progressive accumulation of AQs in both cell-free supernatants and cellular extracts from PAO1, *nfxB*\*, and *nfxB*\**ΔmexD* cultures grown in LB supplemented with 5 mM octanoate.

In agreement with previous findings (65), we found that the intracellular accumulation of AQs and pyocyanin production increase when octanoate is added (Figures 3A, 7C and 7D). However, these increases were similar in all strains, and both the pyocyanin production and the absolute AQs levels reached inside cells were still lower in *nfxB*\* than in PAO1 or *nfXB*\**ΔmexD* (Figures 7A, 7C and 7D). These results indicate that a lower availability of octanoate is not the cause for the impaired QS response displayed by the *nfxB*\* strain.

**Figure 7.**
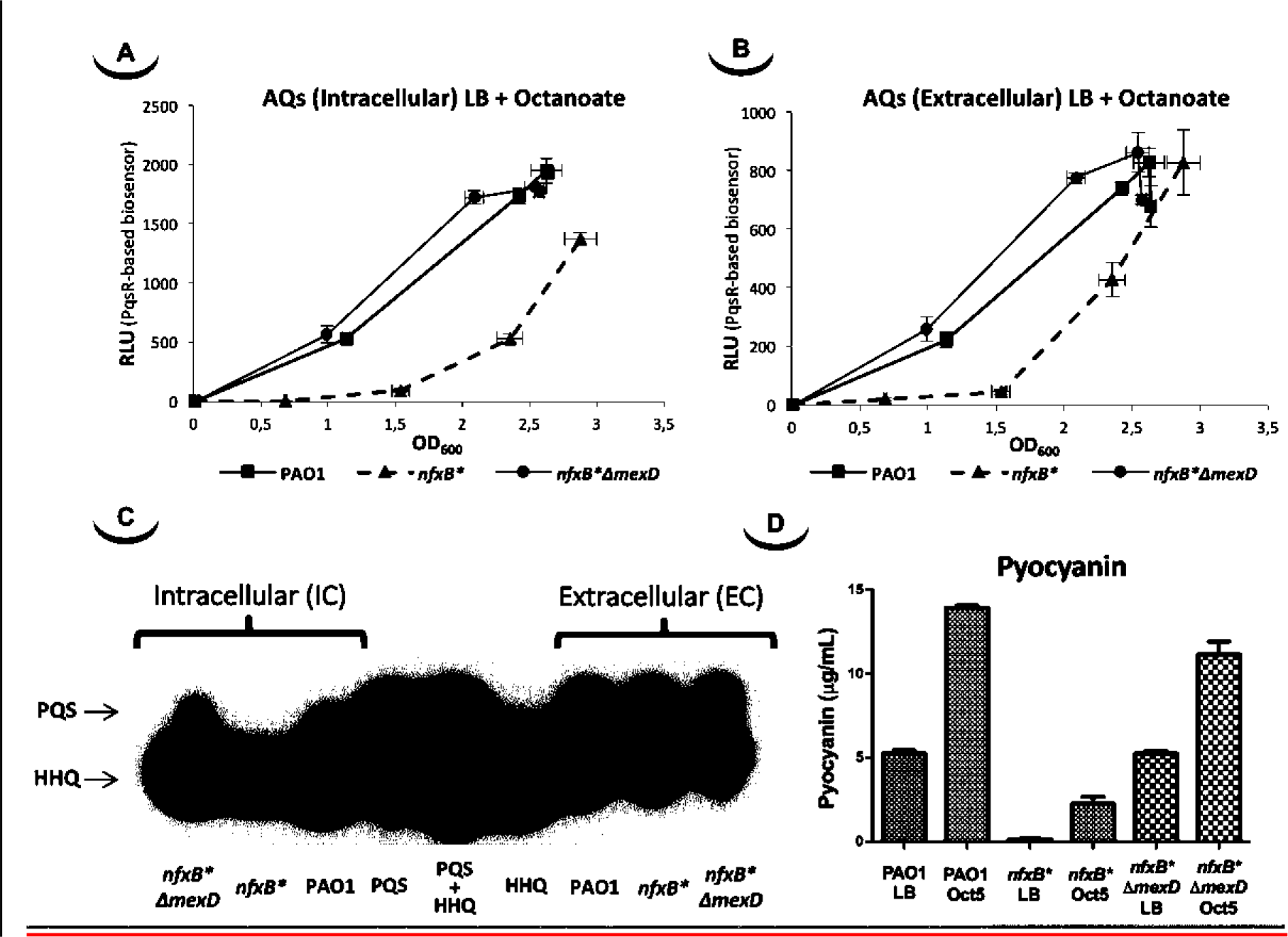
LB supplementation with octanoate increases the AQs production in PAO1, *nfxB*\*, and *nfXB*\**ΔmexD* strains but, in the case of *nfxB*\*, this increase was lower than observed in the other strains, being thus unable to restore the pyocyanin production in the *nfxB*\* strain. To determine the time course production of AQs in PAO1, *nfxB*\*, and *nfXB*\**ΔmexD* strains, we extracted these compounds from both the cells (A) and the cell-free supernatants (B) at different times along the cell cycle (4, 5, 6 and 7 hours post-inoculation). Additionally, the last points of time course extractions were analysed by TLC (C) in order to know the proportion of PQS and HHQ present on each AQs-extracts. For pyocyanin assay (D), the strains were grown in LB medium with or without octanoate (5 mM) along a time lapse of 20 hours and the pyocyanin was extracted with chloroform-based protocol as is described in Materials and Methods. The results show that supplementation of LB with 5 mM octanoate, even allowing *nfxB*\* strain to accumulate similar levels of AQs out of the cells than PAO1 and *nfXB*\**ΔmexD* (B and C), was insufficient to restore neither the intracellular accumulation of PQS and HHQ (A and C) nor pyocyanin production (D). Furthermore, the fact that in a TLC assay (C), the spot corresponding with HHQ present in *nfxB*\* supernatant is slightly higher than that in PAO1 and *nfXB*\**ΔmexD*, together with the clear low intracellular accumulation of HHQ in the *nfxB*\* strain, confirm our hypothesis that MexCD-OprJ is able to extrude HHQ and that is the main reason for the QS-defective response observed in this antibiotic resistant mutant.

It is worth mentioning that, although the *nfxB*\* supernatants exhibit a delay in AQs accumulation in the presence of 5 mM octanoate, the supernatants from all three strains exhibit similar levels when the cultures reach high cell densities (OD_600_ > 2.5) (Figure 7B). In contrast, the intracellular AQs accumulation remains lower in the *nfxB*\* strain (Figure 7A). Further, the analysis by TLC of AQs extracted from the last point of the time course assay showed that, while the intracellular accumulation of PQS and HHQ remained being lower in the case of *nfxB*\*, the extracellular accumulation of these two AQs were similar among PAO1, *nfxB*\* and *njxB*ΔmexD* cultures (Figure 7C). These results further reinforce the hypothesis that MexCD-OprJ is able to extrude HHQ (and likely PQS as well), and confirm that overexpression of this system adversely affects the intracellular accumulation of AQs. This extrusion can be considered the bottleneck that precludes a proficient PQS production as well as the onset of a proper QS response in *nfxB*\*-type mutants.

## Discussion

In the current work, we demonstrate that a *P. aeruginosa nfxB\** mutant, which overexpresses the MexCD-OprJ efflux pump, exhibits an impaired QS response due the extrusion of HHQ. Specifically, this non-physiological extrusion leads to a decrease in expression of the *pqsABCDE* operon responsible for AQs synthesis, which affects AQs-dependent and the PqsE-dependent regulons that comprise the genes involved in swarming motility, and in the production of pyocyanin, rhamnolipids, and proteases among others (50, 59, 66).

The QS response in *P. aeruginosa* consists mainly on the Las, Rhl, and Pqs systems which are dependent on the 3-oxo-C12-HSL, C4-HSL, and PQS/HHQ autoinducers respectively (39). The cross-regulation between these QS-systems is hierarchically understood, with the Las system located at the top, activating the other two QS systems, and followed by the Pqs-dependent activation of the Rhl-system, and by the Rhl-dependent repression of the Pqs-system (39, 67). However, evidence exists that the hierarchy and the relationship between these QS-systems may be modulated depending on environmental conditions and the activity of global regulators as MvaT or RsmA among others (68–72). In addition, recent studies have highlighted the relevant role of the feedback-regulation between Las, Rhl, and Pqs systems as well as the relevance of the PqsE and RhlR regulators (50, 51, 59, 66, 73, 74). Indeed, it has been demonstrated that the expression of approximately 90% of the genes in the AQs-regulon may be regulated through *pqsE* induction (59). Likewise, the non-virulent phenotype prompted by the absence of AQs synthesis may be by-passed through PqsE induction, restoring the full *P. aeruginosa* virulence (51, 59, 66). Further, the regulation of several QS-dependent factors could be redundant. In such a way, the production of elastase, rhamnolipids, or pyocyanin, which are mainly under the control of Las, Rhl, and Pqs systems, respectively, are also regulated by PqsE independently of AQs production (50, 51, 59, 66). In addition, expression of *rhlR* increases upon *pqsE* induction at the same time that some functions of PqsE as a QS-regulator are dependent on RhlR and C4-HSL production, thus establishing a complex feedback regulation loop (59). Even more, exogenous addition of C4-HSL to the cultures may partially complement some of the phenotypes impaired in a *pqsE* mutant, such as pyocyanin production (59, 66, 73).

Given this potential role of PqsE as one of the main QS regulators, we postulate that a reduced production of PQS and HHQ, together with a decreased *pqsE* expression, are the main causes for the lack of QS-response associated to the constitutive overexpression of the MexCD-OprJ efflux system. In this work, we show that expression of QS- regulated genes decreases in an *nfxB*\* antibiotic resistant mutant and that inactivation of the MexCD-OprJ efflux pump in this background restores expression of these genes to wild-type levels (Figure 2). Similar results were obtained with some QS-regulated phenotypes such as the production of elastase, protease IV, pyocyanin, rhamnolipids, and swarming motility (Figure 1), indicating that the alterations in the QS-response displayed by the *nfxB*\* mutant are directly caused by the increased expression and activity of the MexCD-OprJ efflux system. We also demonstrated that loss of function of NfxB leads to an excessive extrusion of HHQ through the overexpressed MexCD-OprJ efflux pump, resulting in a low intracellular accumulation. Expression of *pqsABCDE* during exponential and early stationary growth phases is subjected to a positive feed-back transcriptional regulation under the control of the PqsR-(PQS/HHQ) complex (50). Therefore, the non-physiological HHQ extrusion through MexCD-OprJ may abrogate this positive feed-back regulation and directly cause the decrease in *pqsABCDE-phnAB* expression (Figure 2) and the AQs synthesis impairment (Figure 3) observed in the *nfxB*\* mutant. We also showed that this defective AQs accumulation could not be restored by adding either anthranilate or octanoate, the two PQS/HHQ main precursors (Figures 6 and 7), reinforcing the concept that the main cause for the defective QS-response associated to *nfxB* mutations is an excessive extrusion of HHQ through MexCD-OprJ, and not of metabolic precursors as kynurenine, also extruded by MexCD-OprJ. Additionally, the presence of similar levels of PQS in the supernatants of PAO1, *nfxB*\* and *njxB*ΔmexD* growing in presence of octanoate, together with the absence of this autoinducer signal in the cell-extracts of *nfxB*\* (Figure 7C) suggests that PQS could also be a MexCD-OprJ substrate.

To sum up, here we show that the AQs production is affected by the increased efflux of HHQ by the MexCD-OprJ RND system overexpressed in the *nfxB*\* ciprofloxacin-resistant mutants that are sporadically isolated from ciprofloxacin-treated patients (28). As a consequence, expression of the Pqs-regulon, which also comprises those PqsE-regulated genes in a PQS-independent way (50, 66), is strongly altered in an *nfxB*\* mutant. This alteration may have minor collateral effects on the AHLs-dependent QS systems and is likely the main cause for the low virulence profile observed in antibiotic resistant mutants overproducing MexCD-OprJ.

Moreover, our findings could credit the tightly controlled MexCD-OprJ production with an additional role: signalling at the population, interspecies, and even inter-kingdom levels. It has been shown that, in addition to contributing to a coordinated response of the bacterial population, several QS signal molecules (75–77) are also involved in inter-specific communication networks that modulate the structure and activity of natural microbiomes. The fact that in this work we demonstrated that MexCD-OprJ is able to extrude HHQ, altering the accumulation level of the autoinducer signals produced by *P. aeruginosa*, opens a new perspective over the potential functions of this RND efflux system in the interactions between this opportunistic pathogen and other co-existing species. For example, it has been shown that AQs may function as antimicrobial compounds against *Staphylococcus aureus*, a bacterial species commonly detected together *P. aeruginosa* in polymicrobial infections (78–80). Further, HHQ also is able to induce apoptosis in human mesenchymal stem cells (81), and to impair the production of several factors implicated in the innate immune response affecting the binding of the nuclear factor-кβ to its targets (82). Whether or not MexCD-OprJ overexpression may modulate these interactions remains to be established.

The relationship between MexCD-OprJ and the QS-response demonstrated in this study prompts the speculation about the *nfxB*\* mutants acting as cheaters in *P. aeruginosa* populations. The activation of the QS-response implies an increase in expression of hundreds of genes, and it has been estimated that this consumes approximately 10 % of *P.* aeruginosa metabolic resources (83). Under this panorama, the QS-defective mutants, which commonly emerge in chronic microbial infections and are unable to produce different exoproducts such as siderophores or proteases relevant for nutrients uptake, could be cheaters supported by neighbour bacteria able to produce these QS-dependent factors (84, 85). In this way, the switch-off of QS response in *nfxB*\* mutants would allow them to function as cheaters in mixed population in which they may obtain the benefits brought about by an appropriate QS response carried out by other counterpart bacteria without the cost associated with it. As demonstrated for other systems, it could be predicted that increased abundance of cheaters will produce the collapse of the population and its return to a wild-type situation in a kind of short-sighted evolution (86). Nevertheless, the fact that *nfxB*\* cheaters are resistant to antibiotics has important implications concerning the persistence of antibiotic resistant mutants even in the absence of selection (87).

We propose that MexCD-OprJ should be considered as a new key component in the complex QS-regulation network with a potential role in *P. aeruginosa* intra-species and inter-species signalling. Further, we suggest that a main function of this efflux pump, and of others that reduce the QS response when overexpressed, could be acting as a QS switch of *P. aeruginosa* in response to environmental cues, allowing to switch-off the system in those conditions in which the activation of this energy-expensive global response is not advantageous, despite the population density being high enough to trigger the QS response. Finally, we also propose that MexCD-OprJ overproducer strains might be eventually selected even in the absence of antibiotic selective pressure, acting as antibiotic resistant cheaters in heterogeneous *P. aeruginosa* populations. Since this type of mutants are selected for in infected patients (88) and may keep their pathogenic potential (28), despite their QS defect, this possibility may have potential implications for the treatment of *P. aeruginosa* chronic infections.

## Materials and Methods

### Bacterial strains, plasmids, primers and culture conditions

The *Escherichia coli* and *P. aeruginosa* strains and the plasmids used in this work, are listed in the Table 1. The primers used are listed in the Table 2.

**Table 1.**
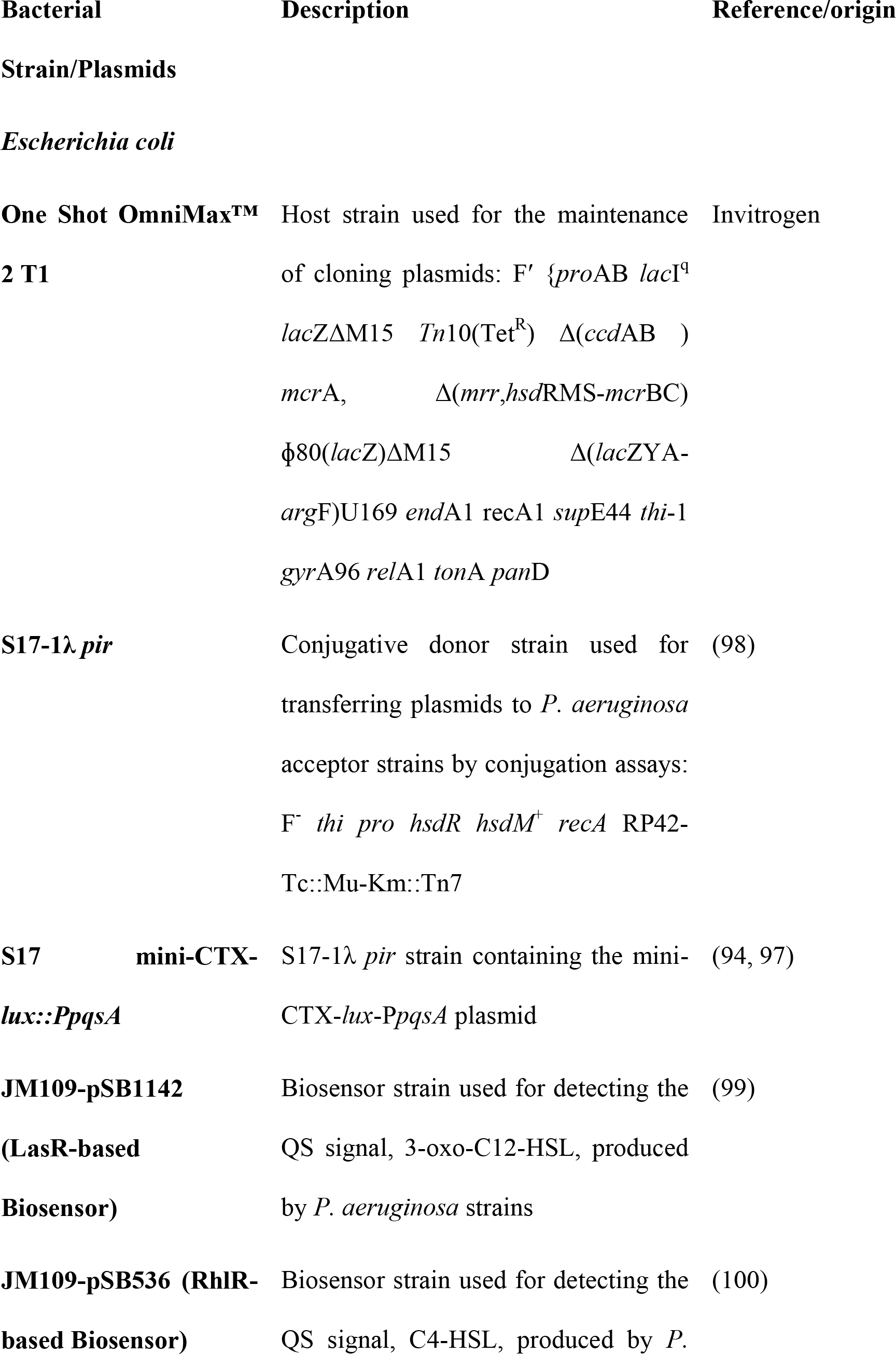
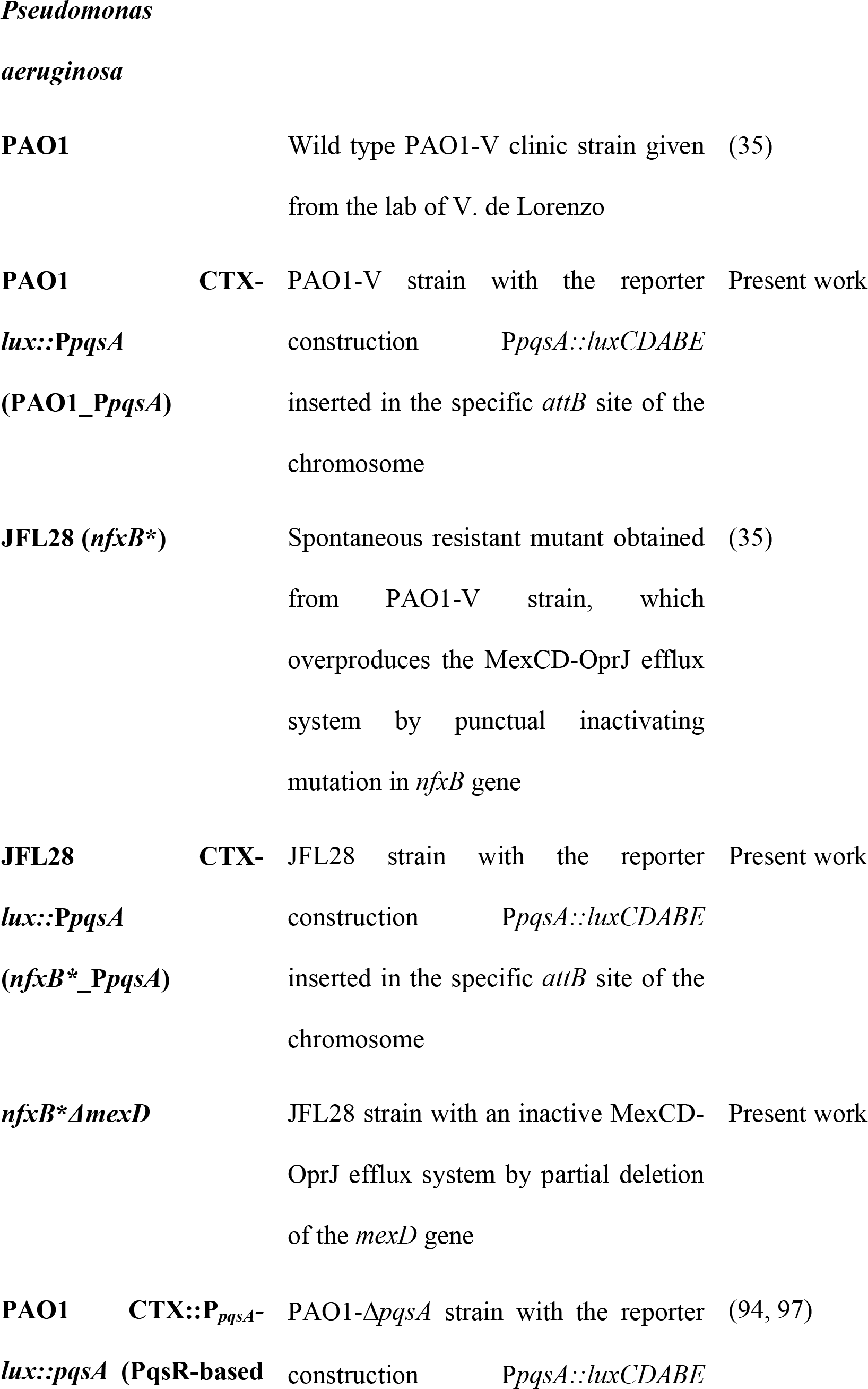
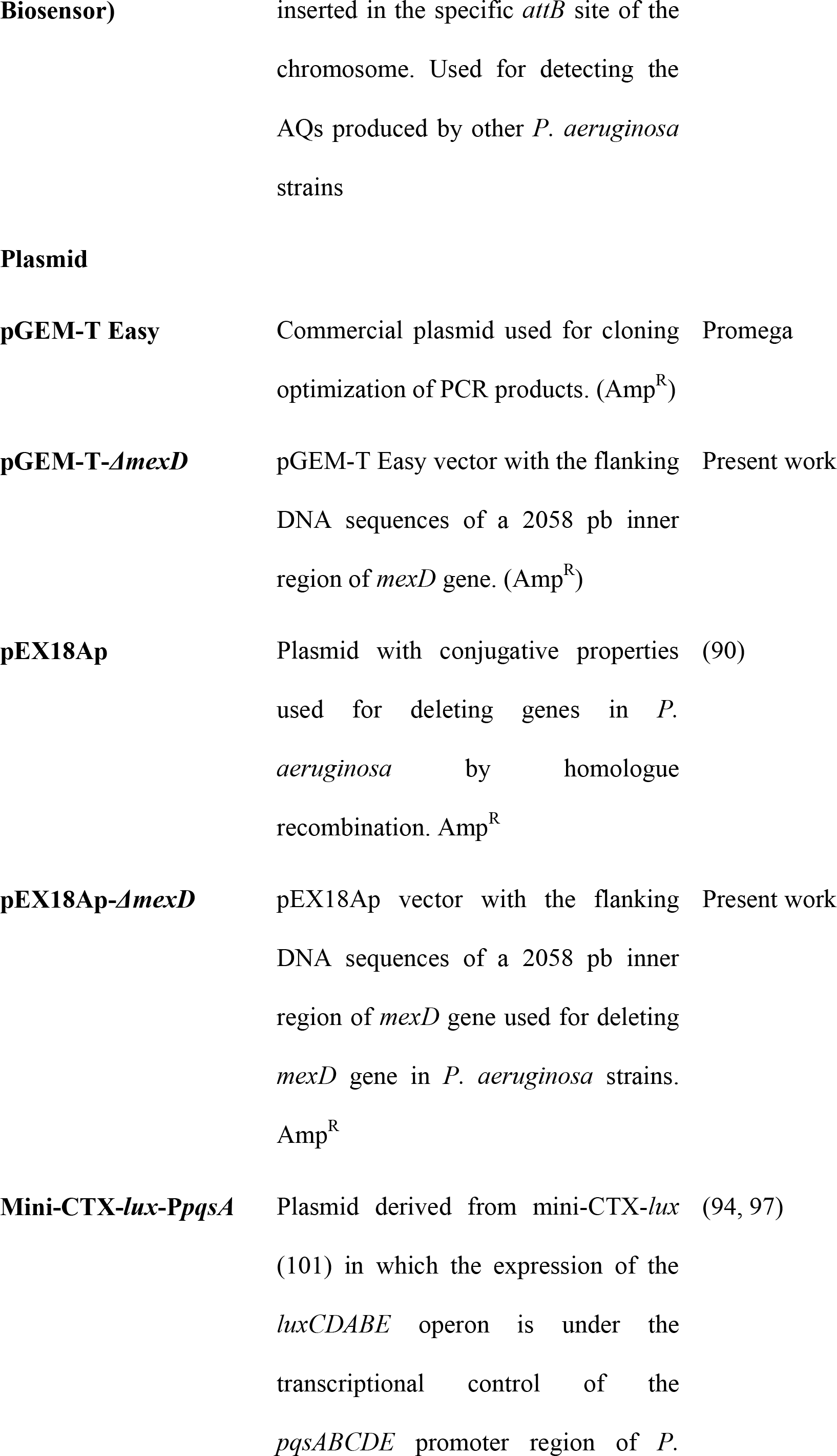
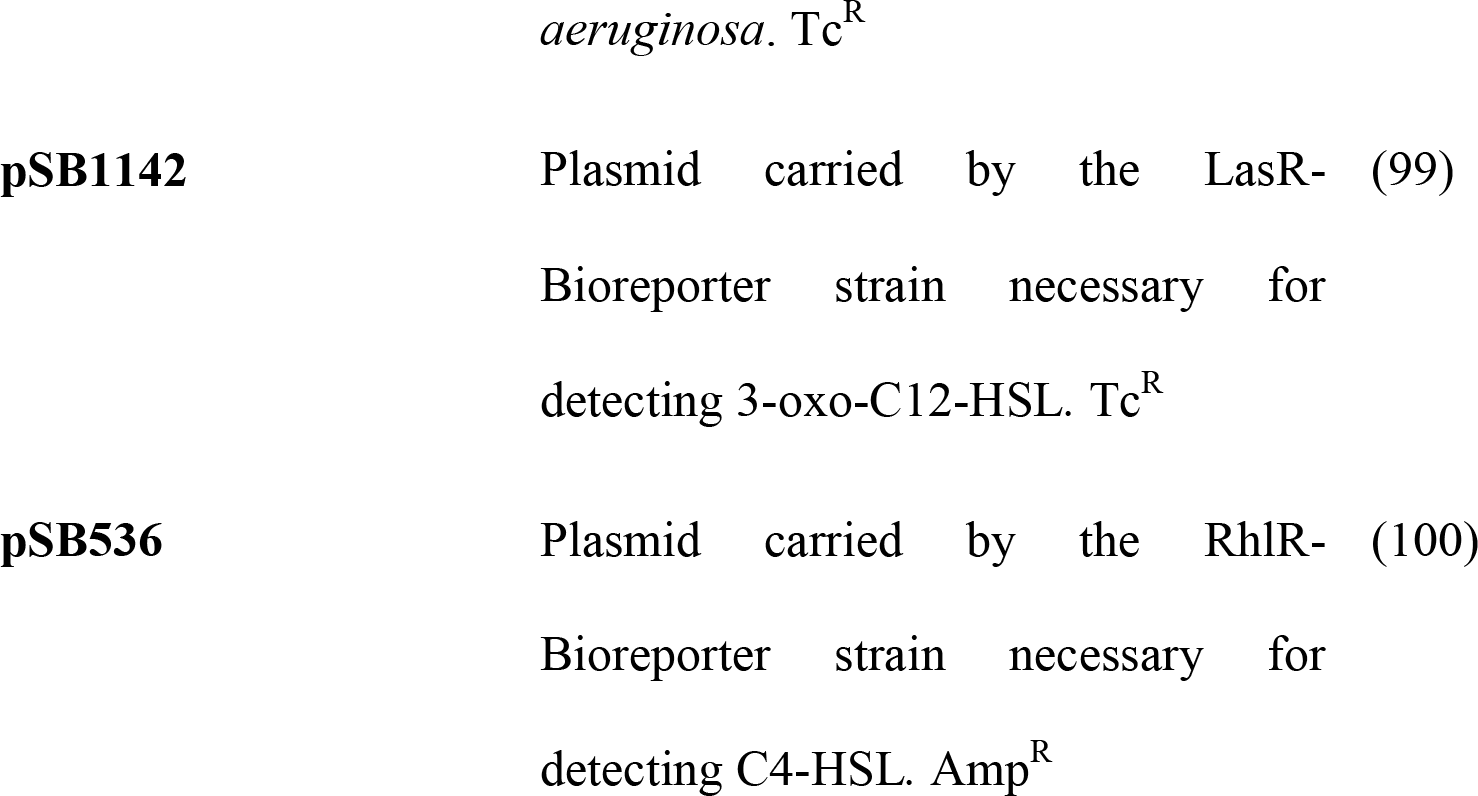
Bacterial strains and plasmids used in the present work.

**Table 2.**
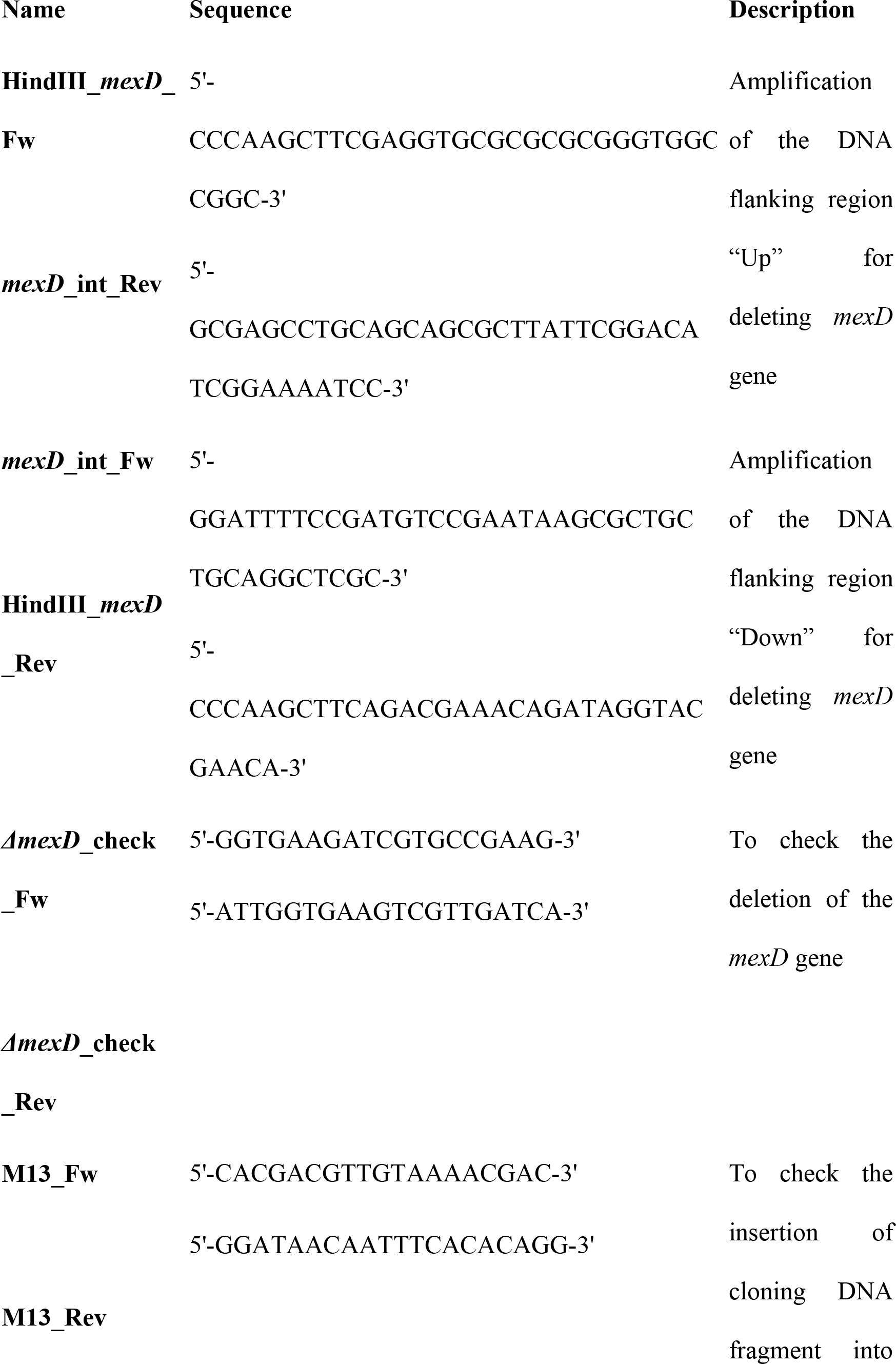
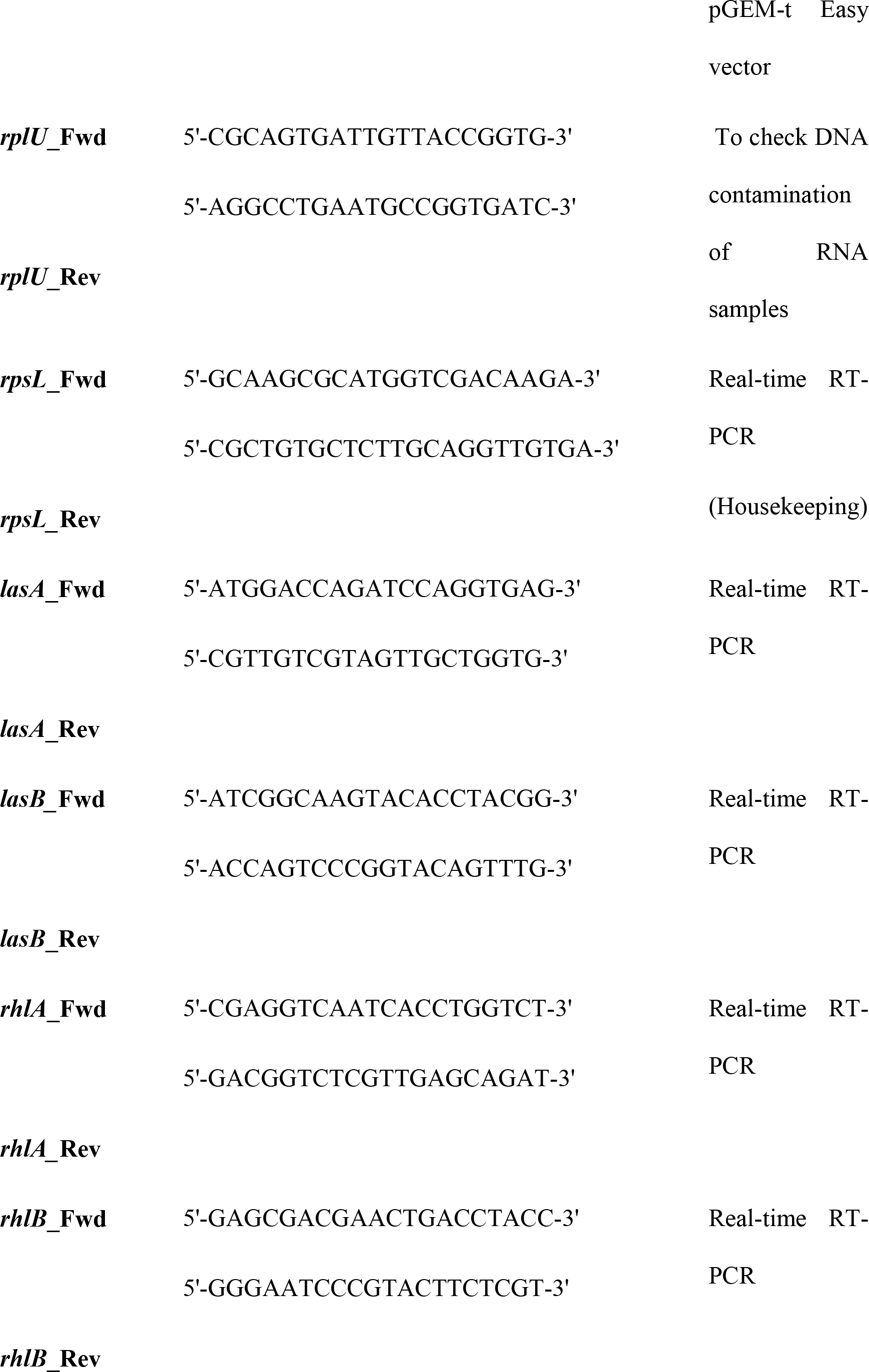
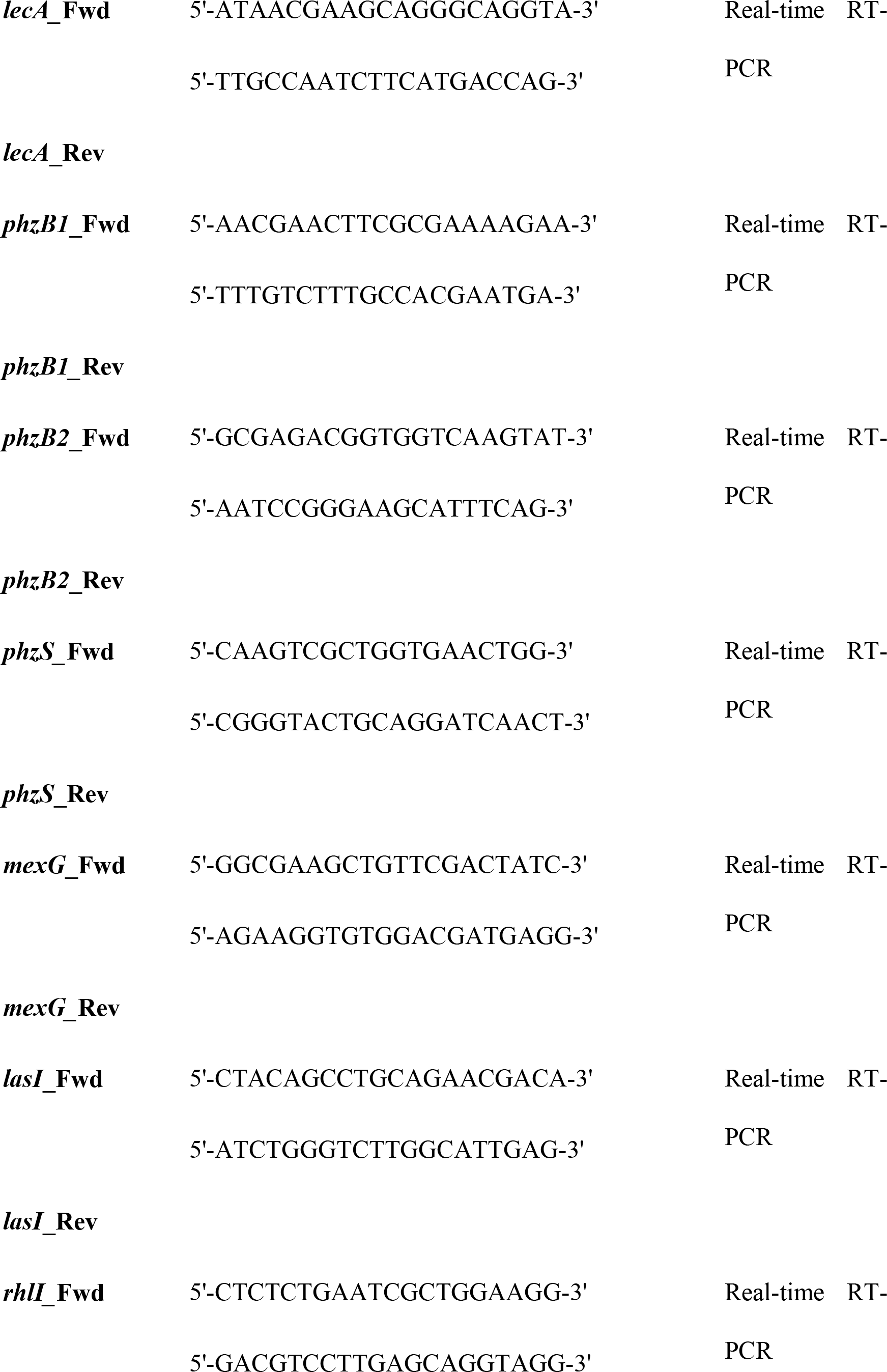
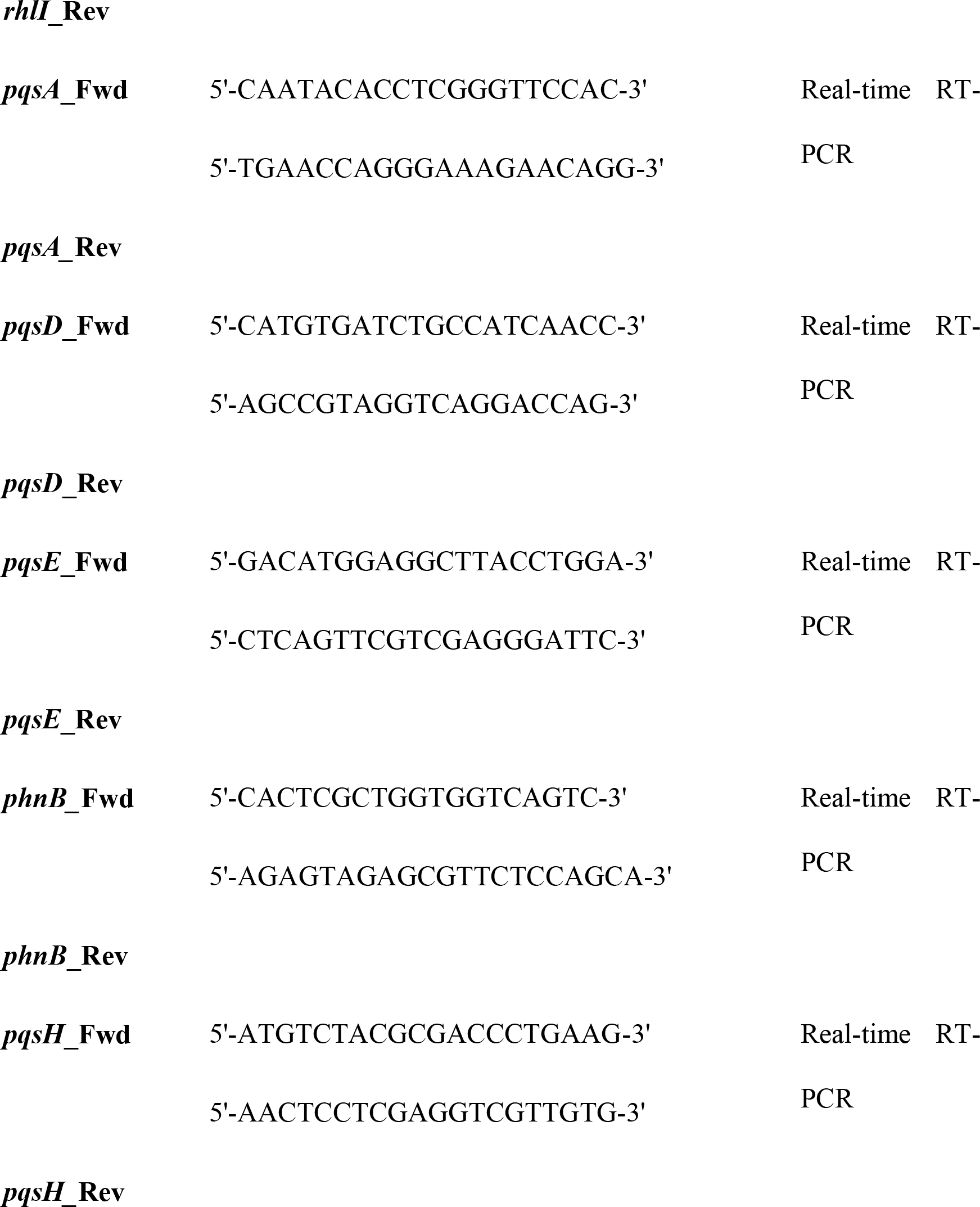
Bacterial strains and plasmids used in the present work.

Unless other conditions are specified, experiments were carried out at 37 °C in 100 ml flasks containing 25 ml of LB broth (Lennox). The *E. coli* strains carrying plasmids with ampicillin (Amp^R^) or tetracycline resistance genes (Tc^R^) were grown in LB medium with 100 μg/ml of ampicillin or 10 μg/ml of tetracycline, respectively. For determining the effect of different carbon sources on *P. aeruginosa* growth, overnight cultures were washed with M63 medium containing MgSO_4_ 1 mM and diluted to an OD_600_ = 0.01 in clear bottom 96-well plates containing 150 μl/well of M63 containing the corresponding carbon source at a final concentration of 10 mM. The growth of each strain was measured at 37 °C using a multi-plate reader.

### Whole genome sequence of the *nfxB*\* strain and generation of a *nfxB*\**ΔmexD* mutant

The *nfxB*\* mutant was fully sequenced at Parque Científico de Madrid using Illumina technology as described (89). Two ≈1000 bp DNA regions adjacent to the fragment of *mexD* to be deleted were amplified by PCR using the primers listed in Table 2. The amplicons were purified and used together for a nested PCR reaction in which a recombinant 2058 bp DNA was generated and cloned into pGEM-t Easy (pGEM-T-*ΔmexD*). *E. coli* OmniMax™ cells were transformed with this plasmid and the sequence of the construction was verified by Sanger sequencing. The fragment was excised using HindIII and subcloned into pEX18Ap. The resulting pEX18Ap-*ΔmexD* construction was incorporated into *E. coli* S17-1λ *pir* by transformation. Introduction of the deleted allele into *P. aeruginosa nfxB*\* was performed by conjugation using S17-1λ *pir* (pEX18Ap-*ΔmexD*) as donor strain as described (90). *mexD* deletion was confirmed by PCR using the primers described in Table 2.

### Analysis of the production of QS-regulated virulence factors

The secretion of elastase and protease IV was measured following the methods described in (55). Rhamnolipids detection was carried out as described (91). Pyocyanin was determined as detailed (92). For the swarming motility assay, O/N cultures were washed with sterile 0.85% NaCl and diluted to an OD_600_ = 1.0. Five-microliters drops were poured on the centre of Petri dishes containing 25 ml of a defined medium (0.5% casamino acids, 0.5% bacto agar, 0.5% glucose, 3.3 mM K_2_HPO_4_ and 3 mM MgSO_4_), which were incubated 16 hours at 37 °C.

### RNA extraction and real-time RT-PCR

RNA was obtained using the “RNeasy mini kit” (QIAGEN) as described (34). After treatment with DNase (34), the presence of DNA contamination was checked by PCR using *rplU* primers. Real-time RT-PCR was performed as described in (34) using the primers listed in Table 2. The experiments were carried out in triplicate. The 2^−ΔΔCt^ method (93) was used for quantifying the results, normalizing the results to the housekeeping gene, *rpsL*.

### Thin Layer Chromatography (TLC) and time course monitoring of QSSMs accumulation

Bacterial O/N cultures were washed with fresh LB medium and diluted to an OD_600_ = 0.01 for subsequent growth. For TLC assays, the QSSMs extractions were carried out as described (94). For time course assays, this protocol was optimized to simultaneous monitoring QSSMs accumulation and cell density. For each extraction time, 1.8 ml aliquots from cultures were centrifuged (7,000x g, 10 minutes at 4 °C). The supernatants were filtered through 0.22 μm pore size membrane and the cellular pellets were resuspended in 1.8 ml of methanol HPLC grade to extract the QSSMs. 900 μl of cell-free supernatants were used to extract the QSSMs by adding 600 μl of acidified ethyl acetate twice. The resulting acidified ethyl acetate extracts were dried and subsequently dissolved in 900 μl of methanol HPLC grade.

AQs were detected by TLC as described (94) using the PAO1 CTX::P_*pqsA*_-*lux* biosensor strain. C4-HSL and 3-oxo-C12-HSL were analysed using the JM109-pSB536 (RhlR-based biosensor) and JM109-pSB1142 (LasR-based biosensor) biosensor strains, respectively (95). The image processing software "ImageJ" was used for densitometry analysis of the light spots.

For time course accumulation assays, flat white 96-well plates with optical bottom were filled with a mix containing 5 μl of sample and 195 μl of a 1/100 dilution of the corresponding O/N biosensor cultures. The experiments were carried out on a multiplate luminometer/spectrophotometer reader. The highest relative light units (RLU = luminescence/OD_600_ ratio) obtained for each biosensor strain and the OD_600_ in which the samples were taken from *P. aeruginosa* cultures were represented.

### Analysis by HPLC-MS of kynurenine and anthranilate accumulation in cell-free supernatants

Bacterial strains were grown in M63 containing succinate (10 mM) and tryptophan (10 mM). After 24 hours at 37° C, the supernatants were filtered through a 0.22 μm pore size membrane and lyophilized. 100 mg of each sample were resuspended in two millilitres of 3 mM ammonium acetate and dissolved in H_2_O/methanol (50/50). The amounts of anthranilate and kynurenine were determined by HPLC-MS at Laboratorio de Cromatografía-SIdI from the Universidad Autónoma de Madrid.

### Insertion of the reporter construction, mini-CTX-*lux-PpqsA*, in the chromosome of *P. aeruginosa* and analysis of *pqsABCDE* expression

The insertion of the mini-CTX-*lux-PpqsA* reporter into the chromosomes of the different *P. aeruginosa* strains was carried out by conjugation as described (96) using a *E. coli* S17-1λ *pir* containing mini-CTX-*lux-PpqsA* (97) as donor strain. The resulting *P. aeruginosa* reporter strains were inoculated in flat white 96-well plates with optical bottom containing 200 μL of LB with or without 4 mM anthranilate at an initial OD_600_ = 0.01. The growth (OD_600_) and the bioluminescence emitted by the *PpqsA::luxCDABE* construction was monitored using a multi-plate reader.

## Acknowledgments

Thanks are given to Diana Palenzuela by her help in the assays of quorum-sensing regulated phenotype.

